# Pytri: A multi-weight detection system for biological entities

**DOI:** 10.1101/2022.05.11.491558

**Authors:** Ali Mehdi, Kieu-Nhi Vu, Felix Lambert

## Abstract

The enumeration of biological entities is a critical part of experimental assays and usually requires large lengths of time. The standard method is to count the entities by hand or with OpenCV-based software, which can lead to inaccurate results. Here, we propose an online platform for biologists consisting of a system with multiple trained machine learning weights to detect various biological entities such as yeast colonies, bacterial colonies, and melanoma clusters. The Pytri model achieved a median relative error rate of 7.56% for bacterial and yeast colonies on Petri dishes, 6.58% for colonies on 96-well plates and 10.28% for melanoma cluster microscopy images. We showcase the application of state-of-the-art deep learning tools in bacterial entity detection, achieving significantly higher accuracy than traditional methods when compared to our base standard manual count.

## Introduction

BIOLOGICAL ASSAYS are a crucial part of molecular biology experiments as analytical tools to determine the concentration or potency of substances^1^. In molecular biology, many assays are conducted by counting formations on plates or alternative media, such as agar plates or a hemocytometer^2^. Counts are usually taken manually despite the time-consuming and tedious nature of the process^3^. Over the years, many non-artificial intelligence-based approaches have been conceived to count biological entities, such as OpenCFU - a colony counting platform, which has gained traction in the biology community, but has failed to garner widespread adoption because it requires the download of specific software and parameters to be adjusted manually^4^. Additionally, there is a lack of advanced mathematics and computational skills needed to create complex software within the field of biology. This renders the task of creating automated systems for the enumeration of biological entities a challenging endeavor for researchers^5^. In recent years, machine learning-based applications such as CFU.Ai have entered the mobile space for colony counting, but the rapid runtime came at the cost of accuracy. These tools either offer overcomplicated menu systems or do not provide functionalities for high-throughput analysis on large-scale datasets. There has also been a lack of a universal platform for biological assays of different types, with most systems focusing on detecting one type of bacterial colonies.

The machine learning applications use computer vision: deriving features and information from images such as biological entity counts. This is accomplished through deep learning and Convolutional Neural Networks (CNNs). The network teaches itself to recognize features through the reduction of images into smaller chunks by applying convolutions - a mathematical function that compresses pixels. Following the training of the network, weights are generated and represent how much each predictor contributes to the detection of the entity in question^6^.

Here, we describe Pytri, a platform that receives images from the user, wherein the user can select the desired weight system to analyze their data: colony-forming unit (CFU) assays and melanoma cluster counting, automated microbial colony picker imagery, and bacterial and yeast colonies. Pytri overcomes the limitations of other platforms, offering cloud computing inference that processes images at a faster rate than local devices, and allows high-throughput analysis for large datasets that would be tedious to do on mobile applications. The multi-weight system also allows a variety of assays to be implemented in one system, which proves useful in industrial research laboratories. The machine learning approach removes the limitations of traditional detection methods, such as OpenCV-based models, where colonies cannot be detected after changes in lighting conditions, image resolution or angles^7^.

## Methods

### Melanoma clusters

We obtained data from experiments involving a melanoma clonogenic assay from the Research Institute of the McGill University Health Centre (RI-MUHC). The images were taken on an EVOS^®^ microscope and showed clusters of different sizes and formations **(Fig. 1D)**. 45 images were collected and had a resolution of 2048 × 1536 px. Assays require the enumeration of melanoma clusters rather than individual melanoma cells. The data was manually counted by a biologist. to set the standard for statistical purposes. The number of clusters ranged from a few to a dozen.

**Figure 1.**
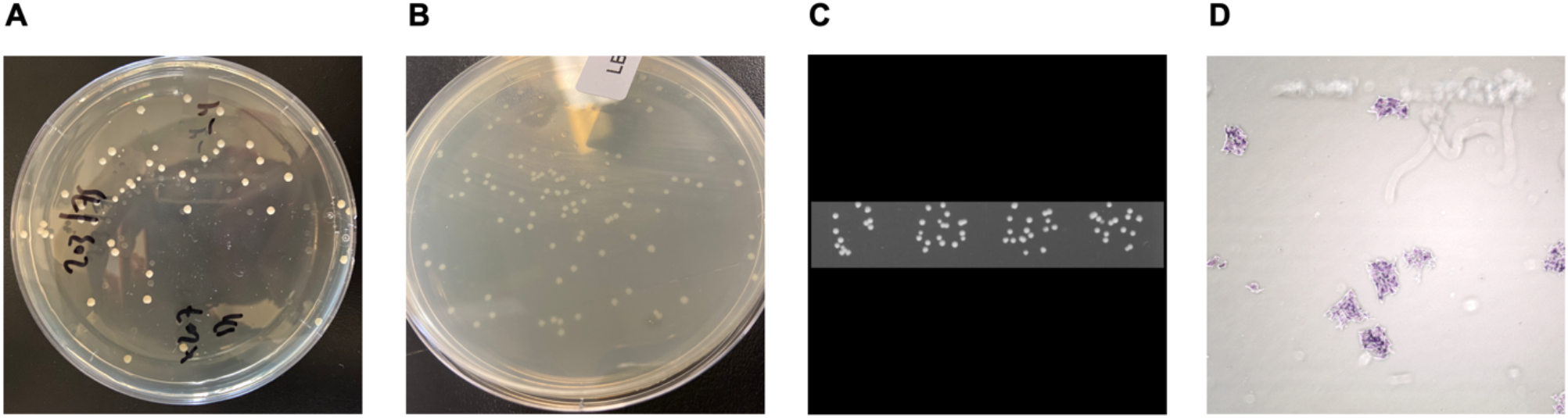
Raw input images used for model training. **(A)** Yeast colonies on a Petri dish grown on agar gel. **(B)** Bacterial colonies on a Petri dish grown on LB media. **(C)** A four-well strip extracted from a 96-well image, with a square black box as background for training. **(D)** A microscopy image of melanoma clusters and individual cells of melanoma.

### *S. cerevisiae* (well colonies)

We obtained data from the Concordia Genome Foundry’s QPix460 camera. Approximately 1100 images were taken at a resolution of 1200×1520 px and with four wells per image **(Fig. 1C)**. The number of colonies per image ranged from about 50 to 150. The QPix460 plates and screens colonies automatically, and images are taken at fixed angles and settings.

### *S. cerevisiae* (plate colonies)

We obtained data from experiments conducted in the Titorenko Lab at Concordia University involving Petri dish plating. The images were taken with a cell phone camera, resulting in resolutions of over 2000×2000 px **(Fig. 1A)**. Initially, we included images with contamination samples to represent both classes (colonies and contaminants) during labeling. However, as more data was added to the dataset, an unequal distribution of classes caused a class imbalance: contaminants (minority class) contained significantly less samples than the colonies (majority class), which resulted in the network failing to detect the entities belonging to the minority class^8^. This was due to the low frequency of contamination relative to the colony count and results in the network omitting the contaminant counts. The final dataset consisted of 125 images. There was variation in the density of the colonies, with some images containing as little as 40-50 and others as much as 500-600 colonies. The images were also taken at various angles and brightness conditions to reflect different situations encountered by researchers^9^. All images had a black bench background. To supplement the dataset, images from the Vogel Lab and the Weber Lab at McGill University were used to add different angles and types of Petri dishes, including rectangular grid dishes.

### *E. coli* (plate colonies)

We obtained data from experiments involving microbial colony growth on LB agar Petri dishes from the Genetics and Cell Biology Laboratory course (BIOL 368) at Concordia University in winter 2021 **(Fig. 1B)**. The images were captured with cell phones, resulting in resolutions of 2000 × 2000 px. 75 images were collected. All images displayed white bacterial colonies similar to yeast colonies. Most images had a black bench background; only a few included a blue bench background. Density did not vary greatly between images (approximately 100 colonies).

### Data pre-processing

For yeast colonies, images with excessive glare or tilted at an extreme angle (60° or lower) were excluded. Remaining images were downsized to a resolution of 416×416 px for faster training. Finally, images of yeast colonies and bacterial colonies were merged due to morphological similarities and formed a dataset of 82 images, referred to as plate colonies **(Table 1)**. To improve the results of the training step, the dataset was augmented by applying a horizontal flip in addition to a variation in saturation of - 25% to 25%. This resulted in 196 images. The dataset was then split into a training set of 172 images, a validation set of 16 images and a testing set of 8 images.

**Table 1.**
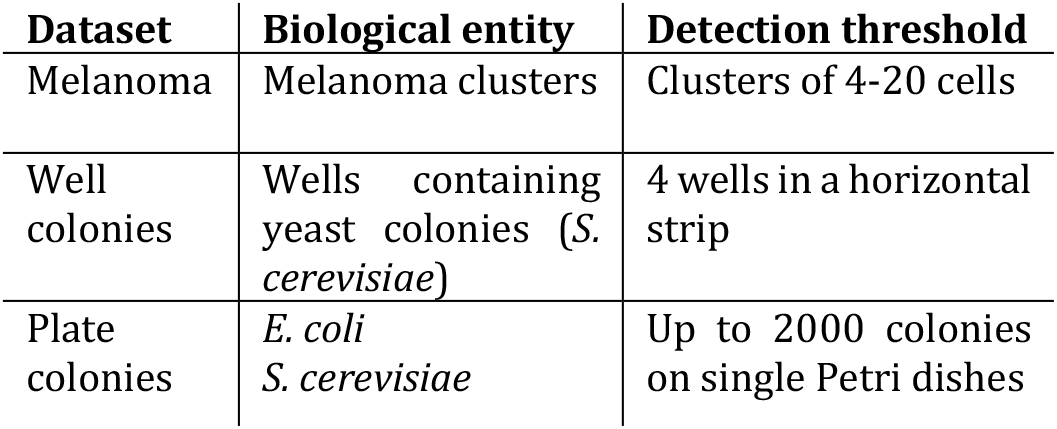
Summary of datasets used in training models and detection.

| Dataset | Biological entity | Detection threshold |
| --- | --- | --- |
| Melanoma | Melanoma clusters | Clusters of 4-20 cells |
| Well colonies | Wells containing yeast colonies (*S. cerevisiae*) | 4 wells in a horizontal strip |
| Plate colonies | *E. coli*<br>*S. cerevisiae* | Up to 2000 colonies on single Petri dishes |

For well colonies, a total of 52 images were selected for training **(Table 1)**. Since images were rectangular, we wrote a script to layer the rectangular slide of colonies over a black square image to avoid squeezing the rectangle into a square and losing accuracy during training. The dataset was augmented by applying both a horizontal and vertical flip, which resulted in 106 images. The dataset was then split into a training set of 90 images, a validation set of 10 images and a testing set of 6 images.

For melanoma cluster microscopy images, the data obtained was limited thus all images were kept. Images were downsized to 1080×1080 due to high picture quality. We included images with many single melanoma cells to strengthen the differentiation between single cells and clusters. The data was augmented by several methods: a 90º rotation, a rotation between −34º and +34 º or the addition of up to 5% noise (to generate conditions of blurred imagery). The final augmented dataset comprised 69 images. It was split into a training set of 60 images, a validation set of 6 images and a testing set of 3 images.

### Model Training

The Ultralytics YOLOv5 model was used for the training phase^10^. YOLOv5 adds built-in image augmentation and adversarial training which modifies the image in the first stage and detects objects in the second stage. Additionally, the YOLO model divides images into grid systems, allowing each grid to detect objects within itself. This becomes crucial with real-time detection and rapid inference^11^.

Google Colab Pro platform was used to train the neural network, providing us with access to high-end GPU (Nvidia Tesla T4). For the model training parameters, the batch size was 16 for all the datasets, the image resolutions were 1080×1080 px, 1024×1024 px and 1018×1018 px for the melanoma, well colonies and plate colonies datasets respectively. The learning rate hyperparameter was modified from 0.01 to 0.005 for all dataset training sessions, which results in better model metrics. The slower learning rate gives a more optimal set of weights, at the cost of a longer training time.

### Benchmarking

We chose the manual count of colonies and clusters as our base standard for comparison. Images were printed onto A4 paper in black and white. For increased accuracy, we used a clicker app to take note of every 10 counts. We also kept track of the time required to count every image with a chronometer.

We compared our manual count of plate colonies - yeast and bacteria merged into one dataset - with the counts of three software: OpenCFU, CFU.Ai. and the YOLOv5-based platform Pytri. For well colonies, the QPix460 picker has an in-built algorithm to identify colonies, but the detection data was not publicly available. For melanoma clusters, no software is currently capable of counting them. Thus, comparison between software was unachievable for well colonies and melanoma clusters, as only Pytri can detect them accurately.

### Statistical tools

The data was analyzed in RStudio version 1.1.456 and with the following additional libraries: ggplot2 (v.3.3.5), tidyverse (v.1.3.1), dplyr (v.1.0.9), ggpubr (v.0.4.0) and car (v.3.0-12). Normality of datasets was assessed using both visual inspection of Q-Q plots and Shapiro-Wilk’s test. Homogeneity of variance was assessed with Levene’s test. Datasets indicating a false assumption of normality or homogeneity of variance were treated with a Kuskal-Wallis test. The results were reported through the p-value’s significance and visually represented in boxplot graphs and scatterplots.

## Results

We evaluated different model architectures and hyperparameters before finalizing our decision on the YOLOv5 model. The YOLOv5 model balanced both accuracy and inference time. Usually, deep learning datasets are large and contain a large sample of images. However, in our application, obtaining a large dataset of quality was difficult due to different factors, such as the time to produce experimental melanoma microscopy imagery, the sensitivity of data with some laboratories and the reliance on university laboratory courses to generate more data from student experiments. We conducted various augmentations to increase the size of our dataset, such as flips, rotations and contrast adjustment.

The plate model was trained using the dataset of 172 images. The training script was run for 150 epochs, resulting in a precision of 0.83. The well colonies model was trained with a dataset of 106 images. The training script was run for 200 epochs, resulting in a precision of 0.91. The melanoma clusters model was trained with a dataset of 69 images. The training script was also run for 150 epochs, resulting in a precision of 0.70 for melanoma cluster detection. The models were halted after plateauing in their precision, while using early stopping. The metric used to assess the model performance is the mAP@0.5 (Mean Average Precision)^12^, which is based on the calculation of the average precision for each class based on model predictions. Taking the average precision for a class with the mean of the individual class precisions. A bounding box is considered a “hit” if the minimum 0.5 IoU is achieved (the Intersection over Union, i.e. the area of overlap over the area of union for two particular bounding boxes)^13^. The melanoma clusters model achieved a mAP@0.5 of 0.67 over 150 epochs **(Fig. 2A)**, the well colonies model achieved a mAP@0.5 of 0.95 over 200 epochs **(Fig. 2B)** and the plate model achieved a mAP@0.5 of 0.79 over 150 epochs **(Fig. 2C)**.

### Comparison of the relative error (against manual counts) of three software in detecting plate colonies

We compared the detection performance of each software to the manual count recorded per plate. There was a statistically significant difference in the relative error between the tested software: Pytri, OpenCFU and CFU.Ai (Levene’s test *p*-value = 2.94 × 10^-05^, Kruskal-Wallis test *p* - value = 9.193 × 10^-11^). Additionally, Pytri was found to have significant differences with both CFU.Ai and OpenCFU (*p* - values = 6.3 × 10^-10^ and 2.8 × 10^-05^ respectively). The error rates of OpenCFU, CFU.Ai and Pytri were also compared to the benchmark manual count of plate colonies **(fig. 3).** The lowest median relative error rate was obtained with Pytri (median=7.56%). There was a 274% increase in accuracy for Pytri compared to CFU.Ai and a 147% increase in accuracy for Pytri compared to OpenCFU.

**Figure 3.**
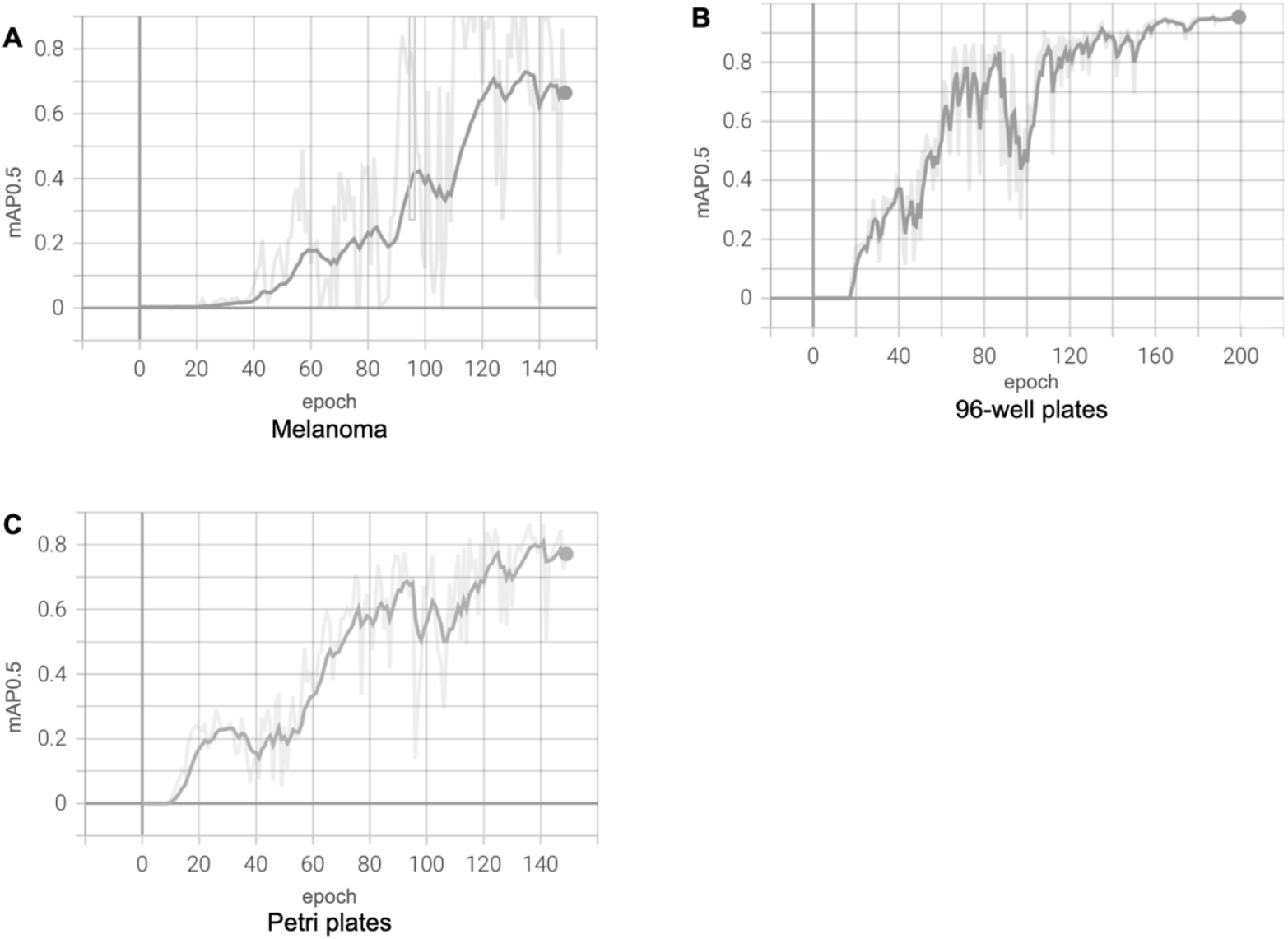
Relative error rates compared to the standard (manual) count of melanoma clusters, colonies in 96-well plates strips and plate colonies (Kruskal-Wallis’ one-way analysis of variance was used for the comparison of relative error rates).

### Pytri’s relative error (against manual counts) in detecting melanoma clusters and well colonies

Data was compared between the manual count of melanoma clusters and well colonies and the count received from Pytri’s output. Due to the lack of software that detects melanoma clusters or well colonies, we could only compare the Pytri model count to the standard count. A Kruskal-Wallis test showed that the Pytri weights in melanoma and well colonies are highly effective at detecting the biological entities (*p*-value = 0.1803, 0.6174 respectively). The Pytri model showed a low relative error rate to manual count for melanoma clusters (median=10.28%) and well colonies (median=6.58%) **(Fig. 3)**.

## Discussion

In this study, we demonstrate the steps required to train multiple detectors in an overarching biological setting. The YOLOv5 convolutional neural network architecture proves to be very advanced at recognizing large numbers of small objects, such as thousands of colonies on Petri dishes while accounting for mirrored colonies on the glass of the dish, which traditional methods fail to recognize. Methods such as OpenCFU depend on the user dedicating a lot of time altering parameters to obtain valid counts, which can be invalidated based on background noise and glare effects. This problem is alleviated in the Pytri model with the inclusion of images that contain glare, reflected colonies, impurities, bubbles in the agar gel and contamination in the training dataset.

In regard to accuracy, the deep learning approach of counting biological entity is more accurate than traditional methods, in both processing time and user-friendliness. For instance, the time required to count colonies tends to increase linearly with the number of colonies on a Petri dish, but this is not the case with colony-counting software **(Fig. 4)**. The interface of OpenCFU is cumbersome and takes longer than solely adjusting confidence thresholds in machine learning approaches, which increases the processing time for data. The QPix460colony picker was unable to detect most colonies. Additionally, even rapid machine learning approaches such as CFU.AI failed to detect colonies as accurately as Pytri, which could be due to increasing inference speed at the cost of accuracy.

**Figure 4.**
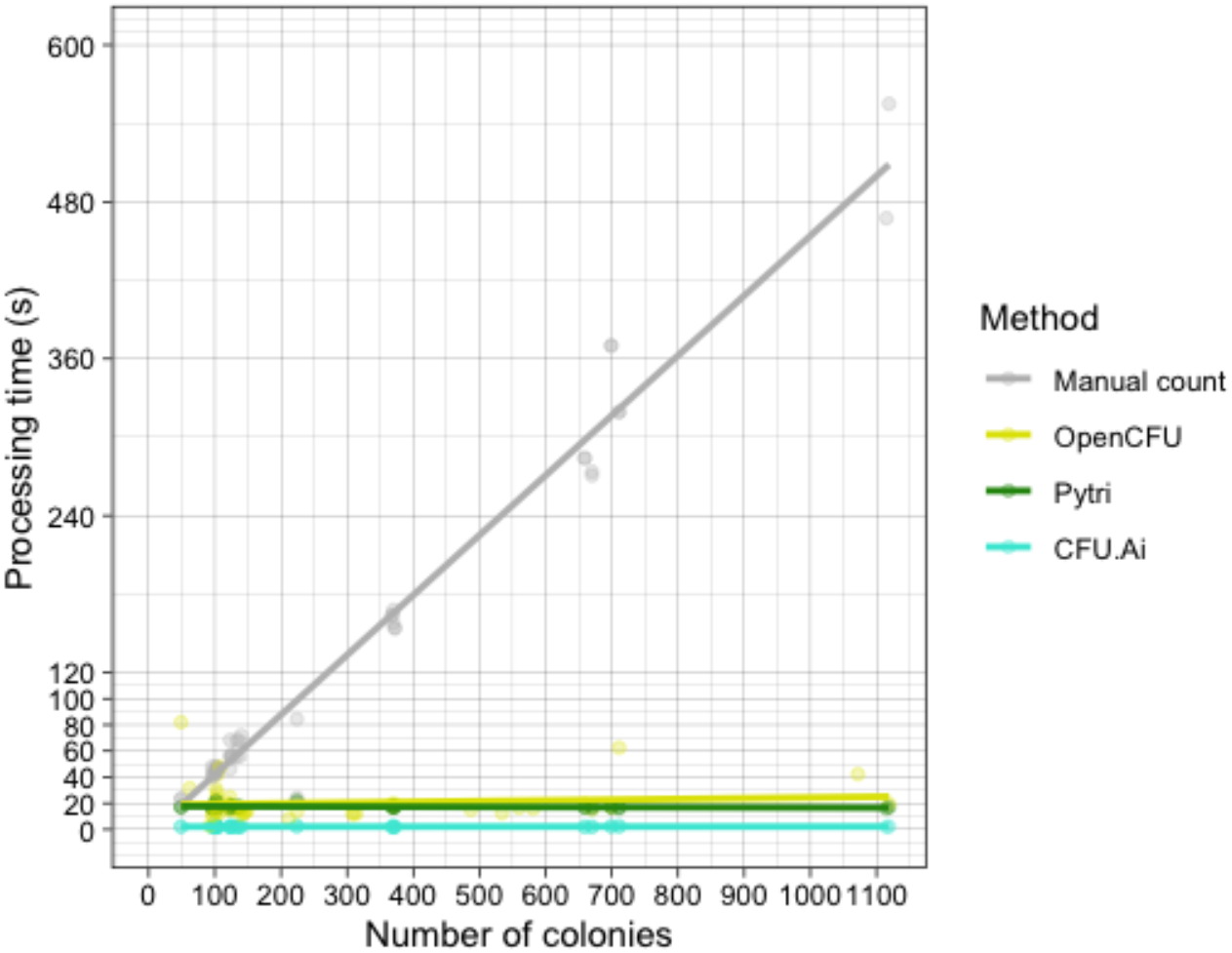
Line plot depicting processing time as a function of the number of colonies detected using manual counting by a biologist, OpenCFU, Pytri and CFU.Ai. The processing time is the sum of the time taken of both manually adjusting the parameters and the computing processing time.

A drawback to the YOLOv5 large model used for Pytri is the slow runtime for large resolutions; this can be alleviated by using YOLOv5 small at the cost of accuracy. The model can be updated with future releases of YOLOv5, such as PP-YOLOv2^14^ and tested with other networks such as Detectron 2^15^.

The training of the model led to lower-than-expected precision and mAP@0.5 results, which could be associated to the heavy colony overlaps in the training data, whereas many colonies overlapped each other in dense formations of 50-150 colonies. The problem was addressed during annotations where clusters were divided into somewhat equal bounding boxes to represent approximate colonies, which could have resulted in the low mAP@0.5 results. However, based on the relative error rates and direct image observations, the machine learning algorithm is robust in accuracy. Training the model with datasets that have more clustered colonies will further improve accuracy and network metrics^16^.

Additionally, it is important to note that error rates are dependent on manual counts, which may not be fully accurate as they are done manually by a scientist. However, in this study, manual counts were established as the standard against which we compared different colony-counting software. Hence, comparison between software is still achievable because the standard count did not vary across software.

Finally, the complexity of building a multi-weight detection platform mainly stems from the subjectivity of researchers when considering biological entity^17^: researchers have different considerations and needs when identifying colonies and clusters (e.g., for the melanoma dataset: the number of cells per cluster, whether to identify individual cells or clusters, etc.). It thus seems necessary to tailor weight systems to specific laboratories, which can take a great amount of time^18^. The generated weights are used to detect biological entities, while the platform returns a colony count with annotated images to the user **(Fig. 5)**. In addition to supporting the four-well strips of QPix460 imagery, single well detection was added by writing a script to crop the image into 96 individual images that are then processed using the plate weight. The results are produced using a 128 grid system with each well being labeled from A1 through L8. Support for additional and more complex assays is being developed, in addition to adding a more user-friendly interface and automation.

**Figure 5.**
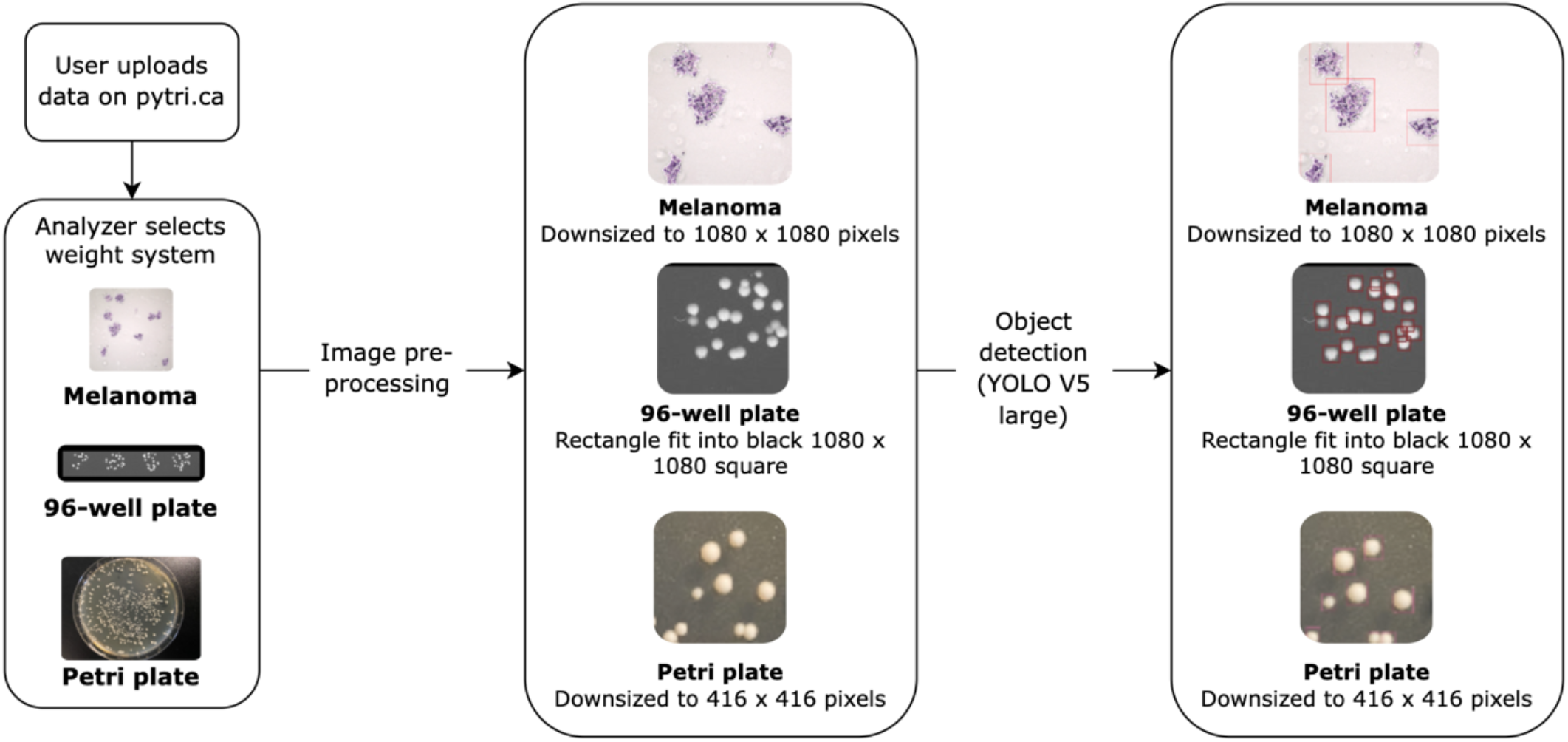
Overview of the processing for the multi-weight detection system. The images are uploaded on the website by the user and then preprocessed according to the selected weight system. The preprocessing is done by downsizing the images and fitting them into square boxes inf they are rectangular. Post-object detection using the YOLO V5 model, the results are returned as a colony count with the images as attachments.

## Conclusion

We have developed a multi-weight pipeline model for accurately enumerating biological entities. This conglomerates biological detection systems into one universal tool capable of analyzing multiple types of biological assays. The pipeline includes weight selection, pre-processing and biological entity detection - all accessible through the online Pytri platform. In the future, the weight system will be expanded to include other types of biological assays, in addition to updating and increasing the accuracy of the models used to generate the weights.

## Availability of data and materials

The code and resultant weight files are proprietary. The data is proprietary to the research labs and from which they were gathered. The data and information are available from the corresponding author A.M. upon reasonable request.

## Competing interests

Ali Mehdi has filed a provisional patent application with the US Patent and Trademark Office and the Canadian Patent Agency related to this work.

## Funding

Research reported in this publication was supported by the Experiential Learning Office, the Dean’s Office, and The Arts and Science Federation of Associations of Concordia University, as well as the Concordia Student Union.

## Acknowledgements

The authors would like to thank Valentina Goanta, Michelle Goanta, Zaki Alasmar, Felipe Perez and Dunia Almelhem for contributing to annotations and other tasks. Special thanks to Michael Shamash for reviewing our work.

The authors would also like to thank the following laboratories, institutes, and people for providing imagery of biological entities used to train the model:

- The Research Institute of the McGill University Health Center
- The Concordia Genome Foundry
- Dr. Titorenko’s lab, Concordia University
- Dr. Madoka Gray-Mitsumune, Concordia University
- The National Institute of Chemical and Pharmacological Research of Bucharest, Romania
- Dr. Jacalyn Vogel’s lab, McGill University
- Dr. Stephanie Crane Weber’s lab, McGill University

## References

1. Bliss, C I. and M Cattell. “Biological assay.” Annual Review of Physiology 5.(1943)

2. Finney, D J.. “The principles of biological assay.” Supplement to the Journal of the Royal Statistical Society 9.(1947): 46–91.

3. Corkidi, G, R Diaz-Uribe, J L. Folch-Mallol and J Nieto-Sotelo. “CO-VASIAM: an image analysis method that allows detection of confluent microbial colonies and colonies of various sizes for automated counting.” Applied and environmental microbiology 64.(1998): 1400–1404.

4. OpenCFU. V3, Quentin Geissmann, 2013

5. Chiang, P J., M J. Tseng, Z S. He and C H. Li. “Automated counting of bacterial colonies by image analysis.” Journal of microbiological methods (2015): 74–82.

6. Albawi, S, T A. Mohammed and S Al-Zawi. (2017, August). n.p.: Understanding of a convolutional neural network. In 2017 international conference on engineering and technology (ICET) (pp. 1–6). Ieee, n.d..

7. Ce, W. (2017, November). n.p.: PCB defect detection USING OPENCV with image subtraction method. In 2017 International Conference on Information Management and Technology (ICIMTech) (pp. 204–209). IEEE, n.d..

8. Japkowicz, N and S Stephen. “The class imbalance problem: A systematic study.” Intelligent data analysis 6.(2002)

9. Shorten, C and T M. Khoshgoftaar. “A survey on image data augmentation for deep learning.” Journal of big data 6.(2019): 1–48.

10. YOLOv5 large. V6.1, Ultralytics, 2022

11. Bochkovskiy, A and C Y. Wang. “Yolov4: Optimal speed and accuracy of object detection.” (2020): 10934.

12. Padilla, R, S L. Netto and E A. Silva. (2020, July). n.p.: A survey on performance metrics for object-detection algorithms. In 2020 international conference on systems, signals and image processing (IWSSIP) (pp. 237–242). IEEE, n.d..

13. Yu, Jiaqian, et al. “Learning Generalized Intersection Over Union for Dense Pixelwise Prediction.” International Conference on Machine Learning. PMLR, 2021.

14. Huang, X, X Wang, W Lv, X Bai, X Long, K Deng and O Yoshie. “PP-YOLOv2: A practical object detector.” (2021): 10419.

15. Pham, Vung, Chau Pham, and Tommy Dang. “Road damage detection and classification with detectron2 and faster r-cnn.” 2020 IEEE International Conference on Big Data (Big Data). IEEE, 2020.

16. Li, X, W Wang, L Wu, S Chen, X Hu, J Li and J Yang. “Generalized focal loss: Learning qualified and distributed bounding boxes for dense object detection.” Advances in Neural Information Processing Systems (2020): 21002–21012.

17. Shamash, M and C F. Maurice. “OnePetri: accelerating common bacteriophage Petri dish assays with computer vision.” PHAGE 2.(2021): 224–231.

18. Ferreira, André C., et al. “Deep learning-based methods for individual recognition in small birds.” Methods in Ecology and Evolution 11.9 (2020): 1072–1085.

