## Supplemental Figure 1 for "Pytri: A multi-weight detection system for biological entities"

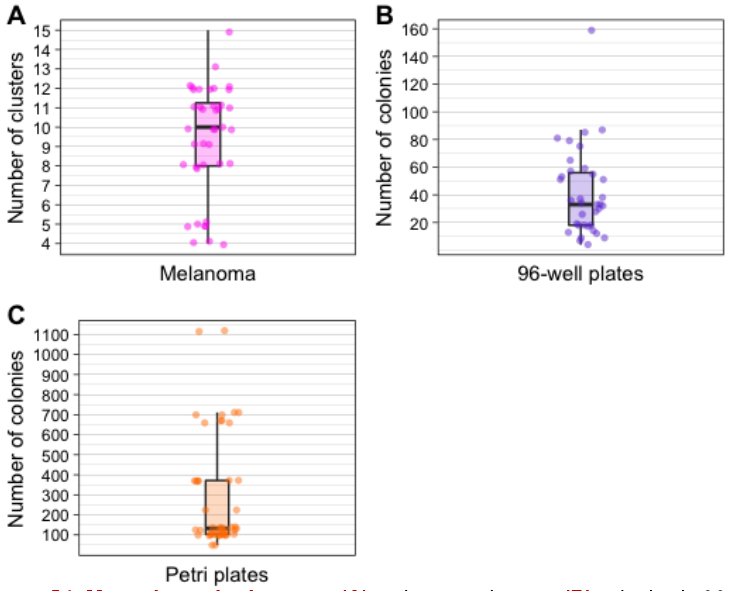


**Figure S1. Manual standard counts. (A)** melanoma clusters; **(B)** colonies in 96-well plates strips and **(C)** yeast and bacterial colonies on Petri plates. Outliers can be seen, representing colony or cluster sizes larger than normal.
